# Individual variation in emergence behaviour and responses to sound exposure in zebrafish

**DOI:** 10.64898/2026.11.15.766749

**Authors:** Fatemeh Alizadeh Lademakhi, Anuradha Bhat, Saeed Shafiei Sabet

## Abstract

Fish respond to sound through behavioural changes, but whether these responses are associated with individual differences in emergence behaviour remains unclear. We examined emergence behaviour across two successive tests and its association with immediate responses to sound in 112 adult zebrafish (*Danio rerio*). Emergence latency was used to classify fish as bold, intermediate, or shy. Latency to enter the upper half of the aquarium was measured before treatment onset, while vertical midline crossings were quantified during 20-s windows immediately before and after sound or ambient-treatment onset. Emergence latency decreased significantly in the second test, and individual latencies were weakly positively correlated across tests. Upper-half latency did not differ significantly between bold and shy fish. Changes in midline crossings differed significantly between treatments, with a greater reduction under sound exposure than under ambient conditions. Exploratory analyses provided insufficient evidence that the association between emergence latency and changes in crossings differed between treatments for either emergence test. These findings demonstrate individual variation and changes in emergence behaviour across successive assessments, alongside an immediate effect of sound on vertical movement. However, they do not establish stable personality differences or emergence latency as a predictor of sound-specific behavioural responses.

## 1 Introduction

Several studies have demonstrated that anthropogenic noise can alter behavioural processes in aquatic organisms among species. In zebrafish, as a model species, controlled sound exposure has been shown to alter swimming speed, swimming height and group cohesion under laboratory conditions. In a separate experiment, fish showed initial behavioural responses but did not avoid the noisier compartment when a quieter alternative was available (Neo et al., 2015a). These findings demonstrate that changes in locomotor behaviour and spatial avoidance need not occur together. Assessing movement before and during exposure can, therefore, help identify immediate behavioural responses that would be missed by considering spatial preference alone.

Behavioural responses to anthropogenic noise can vary among individuals within the same species, and this variation may be obscured when responses are described solely by population-level averages (Harding et al., 2019). This is particularly relevant for behavioural assays in which individuals can differ in exploration, risk-taking, and responses to novel or potentially threatening environments (Conrad et al., 2011; Koets et al., 2025). However, differences observed on a single occasion may reflect both persistent individual tendencies and temporary behavioural states. Repeated observations are therefore needed to evaluate whether individuals retain their relative behavioural differences over time (Bell et al., 2009; Thomson et al., 2020).

Animal personality refers to consistent differences among individuals in behavioural tendencies across repeated observations (Bell et al., 2009). In behavioural ecology, personality research commonly considers dimensions such as boldness, exploration, aggression, and risk-taking. Among fishes, the bold–shy continuum is frequently used to describe individual differences in responses to perceived risk (Conrad et al., 2011). Individual emergence assays can quantify the latency to leave a sheltered compartment (Koets et al., 2025), but interpreting this measure as an indicator of personality requires evidence of consistency across repeated observations (Bell et al., 2009). Consistency within an assay also does not necessarily establish that the same individual differences extend to other behavioural contexts (Bell et al., 2009; Mamede et al., 2024).

Repeated behavioural tests allow changes in average responses to be examined alongside differences among individuals. Individuals may respond differently after previous experience while retaining their relative behavioural tendencies, so behavioural plasticity and consistency can coexist (Dingemanse et al., 2010). In successive emergence tests, a change in mean latency describes an overall shift in emergence behaviour, whereas the association between individual latencies indicates whether fish that emerged earlier in one test also tended to emerge earlier in the next.

Individual differences in boldness may be associated with behavioural and physiological responses to stressors, but emergence latency alone does not establish a stress-coping style (Øverli et al., 2007; Castanheira et al., 2017). In European perch, emergence-based behavioural differences have been linked to stress physiology and immune responses (Gebauer et al., 2023). In zebrafish, Tudorache et al. (2018) found that groups classified by their order of entry into a novel environment differed in their cortisol responses to repeated netting stress. Koets et al. (2025) examined behavioural consistency of zebrafish across time and contexts, as well as the heritability of coping-related traits across generations. These studies motivate examining whether emergence latency is associated with immediate responses to sound, while leaving open whether any such association reflects stable behavioural tendencies or temporary individual states.

Zebrafish (*Danio rerio*) are freshwater fish native to the Indian subcontinent. Field observations in northeastern India found them particularly abundant in vegetated pools with silty substrates and rice paddies adjoining slow-flowing streams (Engeszer et al., 2007). Zebrafish show individual variation in boldness and aggression (Roy & Bhat., 2018), making them a useful model for studying behavioural differences. Their Weberian ossicles mechanically link the swim bladder to the inner ear and contribute to underwater sound detection (Higgs et al., 2002; Popper & Fay., 2011).

Sound exposure can alter zebrafish swimming speed and vertical distribution, with responses varying across exposure conditions and behavioural measures (Neo et al., 2015b; Mohsenpour & Shafiei Sabet., 2021; Wong et al., 2022; Vasconcelos et al., 2023). Here, we focused on vertical midline crossings to quantify movement between the upper and lower halves of the aquarium. Comparing crossings before and immediately after playback onset allowed us to assess whether sound changed the frequency of these transitions. This measure captures one aspect of swimming behaviour, but does not indicate how far fish travelled or how long they remained in either zone.

The present study examined individual variation in emergence behaviour across two successive trials and immediate behavioural responses to experimental sound exposure in adult zebrafish. First, we assessed whether emergence latency changed between trials and whether individual latencies were correlated across trials. Second, we classified fish as bold, intermediate or shy using latency-based thresholds and compared latency to reach the upper half of the aquarium between bold and shy individuals. Third, we compared changes in midline crossings from 20s before exposure to the first 20 s during playback for both ambient and sound treatments. Finally, in an exploratory analysis, we examined whether the association between emergence latency and the change in vertical midline crossings differed between the sound and ambient treatments, considering the first and second emergence latencies separately.

## 2 Materials and Methods

### 2.1. Experimental animals and housing conditions

An initial cohort of 169 adult zebrafish was screened. Of these, 112 individuals completed both emergence tests and were included in the subsequent behavioural analyses. A total of 112 adult zebrafish (*Danio rerio*) of similar age, wild-type coloration (grey and translucent), and mixed sex (approximately 1:1 male:female ratio) were obtained from a commercial aquarium supplier (Astaneh-ye Ashrafiyeh, Iran). Fish were transported to the laboratory and acclimatized for two weeks in a holding tank (86 × 40 × 35 cm; length × width × height) prior to the experiment. Holding conditions were maintained at 25 ± 1 °C water temperature, 14:10 h light:dark photoperiod (lights on at 08:00), and 96 ± 1 lux light intensity. Fish were fed daily with a commercial flake diet throughout the acclimation period; the type and amount of food were kept constant to avoid nutritional influences on behaviour. Prior to behavioural testing, fish were fasted for 24 h to standardise motivational state. Water quality parameters (pH: 8.0 ± 0.2) were monitored daily, and 30 % of the water volume was replaced each day to prevent the accumulation of olfactory cues and organic waste (Wisenden., 2000).

### 2.2. Experimental setup and acoustic characterization

Behavioural trials were conducted in two glass aquaria with identical external dimensions (38 × 25 × 30 cm; length × width × height) and a water depth of 4 cm. The aquaria differed in internal configuration because the second aquarium contained a loudspeaker compartment separated from the test area by a mesh barrier. Each aquarium was divided into two compartments by a fixed opaque partition: a small start chamber (sheltered area) and a larger test area. An emergence door (guillotine type) was fitted into this partition, allowing the fish to move from the start chamber into the test area when raised (emergence door: 2 × 2 cm) (Figure 1 & 3). The start chamber was covered with black plastic to minimise external visual disturbance. Water temperature in the test aquaria was maintained at 27 ± 2 °C, and light intensity was set at 101 ± 1 lux. A plastic barrier separated the experimental area from the observer to minimise human disturbance. The mesh barrier separating the loudspeaker compartment from the test area was positioned 10 cm from the side wall. The barrier prevented fish from physically accessing the loudspeaker.

**Figure 1.**
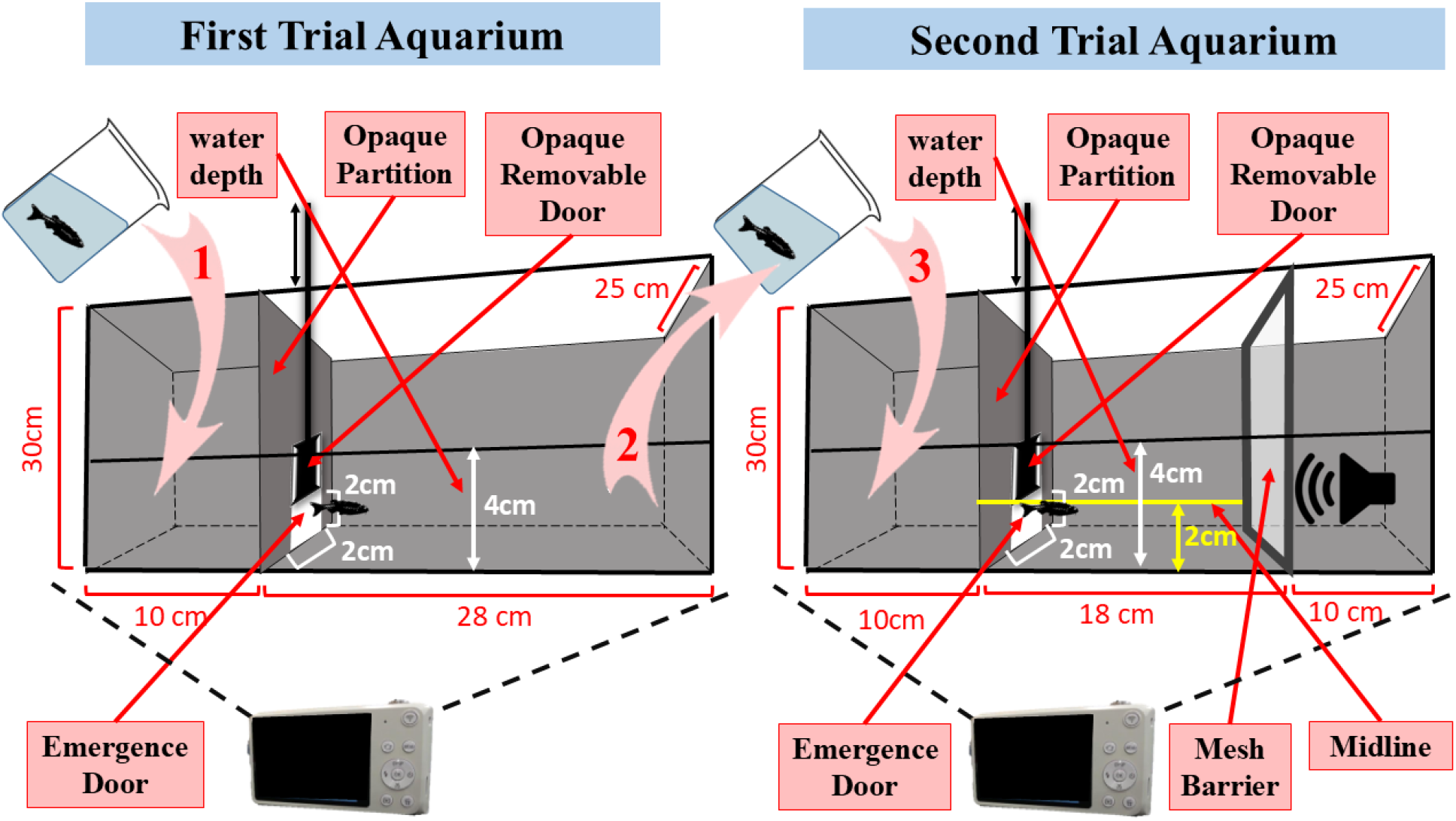
Schematic representation of the two successive emergence tests procedure. (1) A single zebrafish is gently transferred into the start chamber of the first aquarium using a beaker. (2) After emergence from the first aquarium, the same fish is captured again with a beaker. (3) The fish is then transferred to the start chamber of a second aquarium with the same external dimensions as the first for the second emergence trial.

Broadband white noise was generated using Audacity (version 2.3.1; see Mohsenpour & Shafiei Sabet, 2021) and played through a JBL GO2 Bluetooth speaker enclosed in a waterproof vacuum bag and suspended in the water column near the bottom of the test aquarium. Two treatments were used: (i) a continuous sound stimulus (400–2000 Hz, mean sound pressure level: 104.7 ± 1 dB re 1 µPa) and (ii) an ambient (96.4 ± 1 dB re 1 µPa ambient noise). The frequency range was chosen to match the hearing sensitivity of zebrafish (Higgs et al., 2002) and to overlap with anthropogenic noise spectra commonly found in laboratory and aquaculture settings (Slabbekoorn et al., 2010). Underwater sound was recorded using an AS-1 hydrophone (Aquarian Scientific, USA), connected to a PA-4 hydrophone preamplifier (Aquarian Audio, USA) and a TASCAM DR-100MKII digital recorder. This recording configuration was also used in our previous zebrafish research (Mohsenpour & Shafiei Sabet., 2021). The manufacturer specifies a nominal receiving sensitivity of −208 dB re 1 V/µPa for the AS-1 hydrophone. Recordings obtained under ambient and playback conditions were analyzed using custom R scripts to characterize sound pressure in the experimental aquarium. Only sound pressure was measured; particle motion was not quantified. (Shafiei Sabet & Alizadeh Lademakhi., 2022). The acoustic field within the aquarium was characterised to confirm that the sound stimulus was consistently delivered.

### 2.3. Emergence behaviour: two successive

Emergence behaviour was assessed using two successive tests adapted from Tudorache et al. (2018). Each fish was tested individually. At the start of each trial, a single zebrafish was gently captured with a beaker and placed into the start chamber of the first test aquarium under ambient laboratory sound conditions with no playbacks. After a 15 min habituation period, the opaque door was removed, allowing the fish to voluntarily exit the sheltered area and enter the novel, potentially risky environment. The latency to emerge (time taken to leave the start chamber, in seconds) was recorded for each individual. An initial stock of 169 adult zebrafish was screened; 112 individuals successfully completed both successive emergence trials and were retained for the subsequent behavioural analyses, whereas the remaining fish that failed to emerge within 20 min during the first and/or second trial were excluded from the experiment.

Immediately after the first emergence trial, the same fish was transferred to the start chamber of the second aquarium, and the entire procedure was repeated (15 min habituation, door removal, and recording of emergence latency). We therefore did all acoustic playbacks and relevant sound related behavioural assessments in the second tank. This design allowed us to compare emergence latencies across two successive tests and examine whether individuals that emerged earlier in the first test also tended to emerge earlier in the second (Figures 1 and 2). The correlation between latencies was used to assess this association, rather than to establish stable personality traits. (Tudorache et al., 2018; Roy & Bhat., 2018).

**Figure 2.**
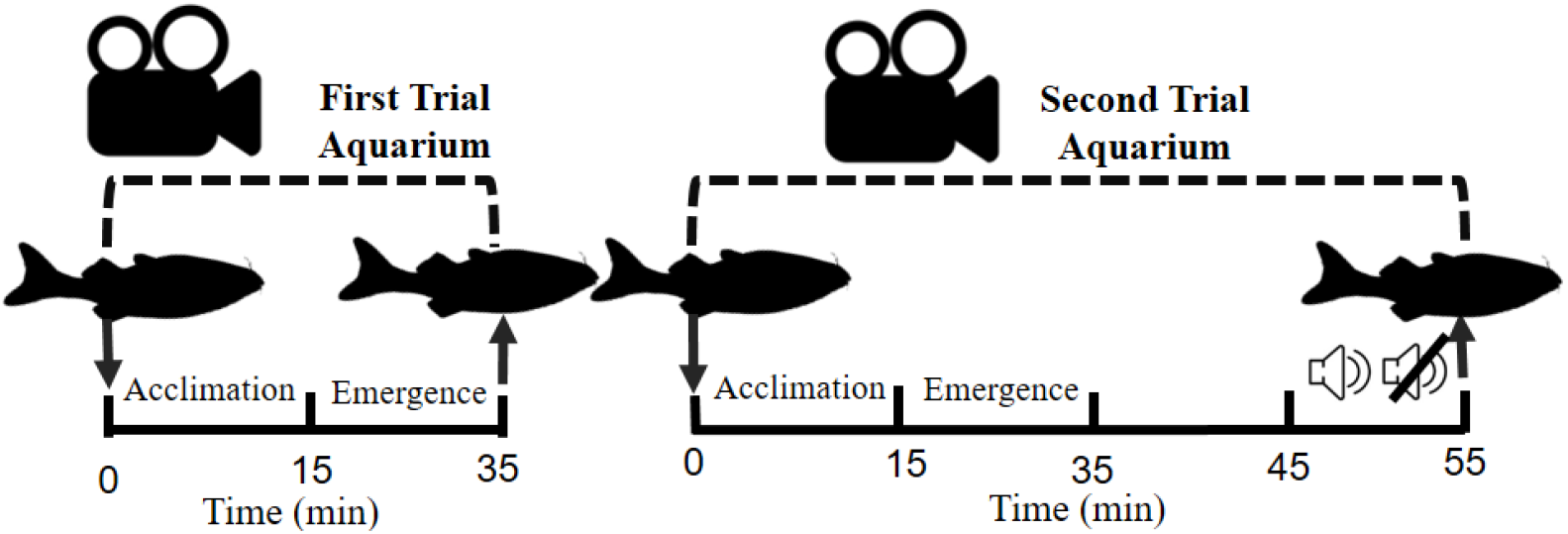
Experimental timeline of the two successive emergence tests and sound exposure procedure. Times are presented as maximum values (e.g., a maximum of 20 min was allowed for emergence; individuals that did not emerge within this period were excluded from further analysis). At each stage, the exact emergence latency was recorded for each fish. Following the second emergence and a 10- min interval, the fish was exposed to 10 min of sound or ambient treatment, during which behavioural parameters were recorded. All times in the figure are in minutes.

**Figure 3.**
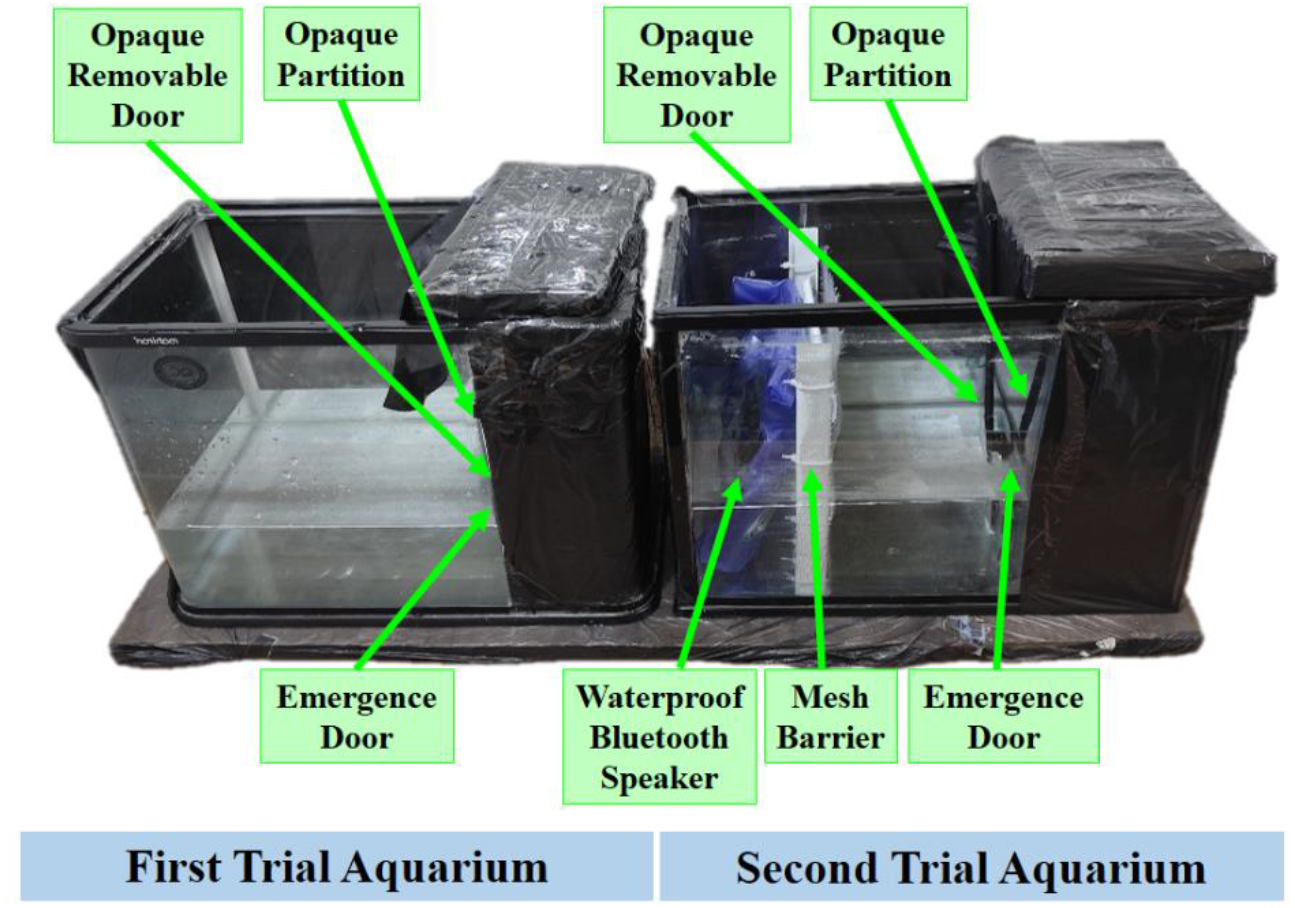
Photograph of the experimental setup showing the two aquaria with identical external dimensions used for the two successive emergence tests. The left aquarium was used for the first emergence trial, and the right aquarium was used for the second emergence trial. Each aquarium was divided into a start chamber and a test area, separated by an opaque partition with a removable door. The second aquarium additionally contained a loudspeaker compartment separated from the test area by a mesh barrier.

### 2.4. Sound exposure and behavioural measurements

Following emergence from the start chamber in the second aquarium, each fish remained in the test area for 10 min before the start of the assigned sound or ambient treatment. The treatment period lasted 10 min. Midline crossings were quantified during the 20 s immediately preceding treatment onset and the first 20 s following onset (See Figure 1). For the ambient treatment, the same observation windows were defined relative to the corresponding time point, without initiating sound playback. Treatments were randomly assigned to minimize experimental error. The loudspeaker, waterproof enclosure and mesh barrier remained in the same positions during both treatments. The loudspeaker remained switched on only in sound treatments. During ambient treatment trials, the loudspeaker was switched off with no playback, whereas during sound treatment trials, it played the experimental acoustic stimulus. Handling procedures and observation timings were otherwise identical between treatments. Behavioural responses were recorded using a digital video camera Samsung WB30F, 50 fps) positioned in front of the aquaria. Videos were converted to 5 frames s⁻ ¹ using Xilisoft Video Converter and analyzed with Logger Pro (version 3.6.3) to track swimming trajectories and calculate behavioural parameters.

The water column was 4 cm deep and was divided analytically into two equal vertical zones, with the horizontal midline located 2 cm above the bottom. This boundary provided a consistent operational definition for measuring latency to enter the upper half and the number of transitions between the two halves. These variables were used to describe vertical movement within the emergence-test arena. Because depth and surface-related cues can influence zebrafish behaviour, these measurements were not treated as equivalent to validated anxiety endpoints from the novel tank diving test (Blaser & Goldsteinholm., 2012; Blaser & Rosemberg., 2012). The following behavioural parameters were quantified for each individual:

i. Emergence latency (ET): The time (s) required for an individual to leave the sheltered start chamber and enter the main test area was recorded during both successive emergence trials (ET1 and ET2). In the second emergence trial, latency was recorded during the initial part of the trial, before the onset of either the sound or ambient treatment. These measurements were used to assess consistency of emergence behaviour across repeated trials and to classify individuals along the Bold– Intermediate–Shy continuum.
ii. Latency to reach the upper half was defined as the elapsed time from emergence into the test area until the fish’s entire body first entered the upper half of the water column, above the horizontal midline located 2 cm above the bottom. This endpoint represented the initiation of upper-water-column use; “free swimming” was not scored as a separate behavioural event. Individuals that did not reach the upper half within 10 min were excluded from the analysis of this parameter. This measure was used to examine whether emergence-based behavioural phenotype was associated with subsequent exploratory behaviour. This parameter was measured after the second emergence test, during the 10 min interval before the onset of the sound or ambient treatment. Of the 112 fish, two did not reach the upper half of the aquarium within this 10 min period and were excluded from the analysis of this parameter.
iii. Midline crossings were defined as complete transitions of the fish’s entire body across the horizontal boundary separating the upper and lower halves of the water column. Each complete transition in either direction was counted once; therefore, a complete crossing followed by a complete return crossing contributed two crossings. Partial crossings, in which part of the body remained on the original side of the boundary, were not counted. Rapid successive transitions were counted separately whenever the entire body crossed the boundary. Crossings were scored during the 20 s immediately before treatment onset and the first 20 s after onset, using the corresponding time point for the ambient treatment. Although behaviour was video-recorded throughout the 10-min treatment period, crossing counts were analyzed during equal 20-s windows immediately before and after treatment onset to characterize immediate changes in vertical movement. Corresponding windows were analyzed in the ambient treatment. This sampling approach addressed the initial response to playback onset and did not assess whether behavioural changes persisted throughout the treatment period.

### 2.5. Emergence-based behavioural classification

A latency-based thresholding approach was used to classify zebrafish into Bold, Intermediate, and Shy behavioural phenotypes separately for the first (E1) and second (E2) emergence tests. For each test, the mean emergence latency and standard deviation (SD) were calculated across the 112 fish. Individuals with emergence latencies below the mean − 0.5 SD were classified as Bold, whereas those with latencies above the mean + 0.5 SD were classified as Shy. Individuals with latencies within ± 0.5 SD of the mean were classified as Intermediate. This procedure allowed the distribution of behavioural phenotypes to be determined independently for each emergence test.

### 2.6. Statistical analysis

Statistical analyses were performed using SPSS (version 26) and R (version 3.6.3). For paired comparisons, the distributions of within-individual differences were assessed using Q–Q plots and Shapiro–Wilk tests. These differences comprised second minus first emergence latency and during-exposure minus before-exposure crossing counts within each treatment. Normality tests were interpreted alongside the graphical assessments, sample sizes and discrete nature of the crossing counts. As a sensitivity analysis, percentile bootstrap 95% confidence intervals for the mean differences were calculated using 20,000 resamples. Fish-level differences were resampled within each comparison, preserving the paired structure of the observations. Emergence latencies were compared between the first and second tests using a paired-samples t-test. Pearson’s correlation coefficient was calculated to assess the association between individual emergence latencies across the two tests; this analysis was not treated as a variance-component estimate of behavioural repeatability. Latency to reach the upper half of the aquarium was compared between Bold and Shy individuals classified using the first emergence test using an independent-samples t-test. Intermediate individuals were excluded from this comparison.

As an exploratory, complementary analysis, the comparison of upper-half latency between bold and shy fish was repeated using the E2-based classification and independent-samples t-test. This analysis used the same upper-half latency measurements, recorded after E2 and before acoustic-treatment onset, and excluded intermediate individuals and fish that did not reach the upper half within 10 min.

For midline crossings, differences between the before-exposure and during-exposure periods were assessed using paired-samples t-tests separately for the sound and ambient treatments. A change score was calculated for each fish as the number of crossings during exposure minus the number before exposure (Δ = during − before). These change scores were compared between treatments using an independent-samples t-test to test directly whether the change in vertical movement differed between sound and ambient conditions.

An exploratory analysis, developed following initial examination of the results and not specified a priori, examined whether the association between emergence latency and changes in vertical midline crossings differed between treatments. Separate linear models were fitted for the first and second emergence latencies, with the change score as the response variable and emergence latency, treatment, and their interaction as predictors. Emergence latency was scaled in units of 100 s to facilitate interpretation of the regression coefficients. Heteroscedasticity-robust HC3 standard errors were used to calculate 95% confidence intervals and conduct two-sided t-tests with residual degrees of freedom. The interaction coefficient represented the difference in slopes between the sound and ambient treatments. Both models used the same individuals and behavioural change scores, with a different emergence-latency predictor in each model. All tests were two-sided, with statistical significance assessed at α = 0.05. Descriptive data are presented as means ± standard error (SE), unless otherwise stated.

#### Ethical approval

The study was approved by the Research Ethics Committees of the University of Guilan (IR.GUILAN.REC.1403.093) and conducted in accordance with the ASAB guidelines for animal research (2023). Fish were handled gently, and handling was limited to the transfers required by the experimental procedure to minimize disturbance. No fish were euthanized, and no mortality occurred during the experiment.

## 3 Results

### 3.1. Distribution of bold, intermediate, and shy phenotypes across repeated emergence tests

To characterize individual behavioural variation, zebrafish were classified into Bold, Intermediate, and Shy phenotypes separately for the first (E1) and second (E2) emergence tests (Figure 4). Individuals with emergence latencies below the mean − 0.5 SD were classified as Bold, those with latencies above the mean + 0.5 SD as Shy, and those falling between these thresholds as Intermediate. In E1, 43 fish (38.4%) were classified as Bold, 40 (35.7%) as Intermediate, and 29 (25.9%) as Shy. In E2, 44 fish (39.3%) were classified as Bold, 44 (39.3%) as Intermediate, and 24 (21.4%) as Shy. Overall, the proportional distribution of behavioural phenotypes was broadly similar between the two emergence tests. The proportion of Shy individuals was lower in E2 than in E1, whereas the proportions of Bold and Intermediate individuals were slightly higher. These results describe the distribution of emergence-based behavioural phenotypes in each test, without implying that individual fish necessarily retained the same classification across tests.

**Figure 4.**
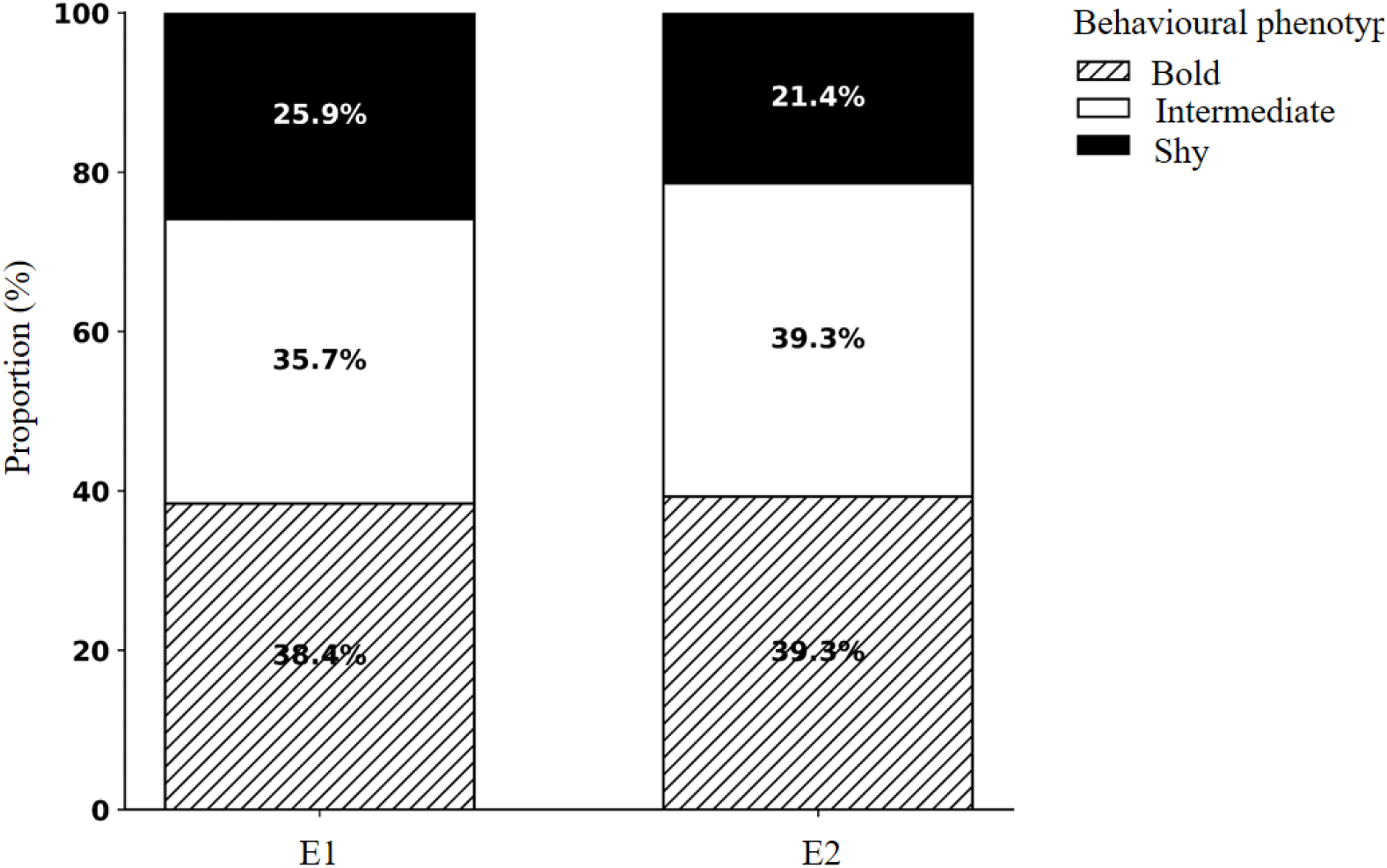
Proportional distribution of behavioural phenotypes across repeated emergence tests. Stacked bars show the proportion (%) of zebrafish classified as Bold, Intermediate, or Shy in the first emergence test (E1) and second emergence test (E2) using latency-based thresholding. Each test included 112 fish. Bold, Intermediate, and Shy phenotypes are represented by hatched white, plain white, and black segments, respectively. Percentages within the bars indicate the proportion of fish assigned to each phenotype.

### 3.2. Changes in emergence latency between repeated emergence tests

Individual emergence latency showed considerable variation among zebrafish and generally decreased from the first emergence test (E1) to the second emergence test (E2) (Figure 5). The mean emergence latency decreased from 381.04 ± 27.98 s in E1 to 153.27 ± 15.37 s in E2. Individual trajectories showed that most fish exhibited a reduction in emergence latency during the second test, although the magnitude of this change varied substantially among individuals. (Figure 5).

**Figure 5.**
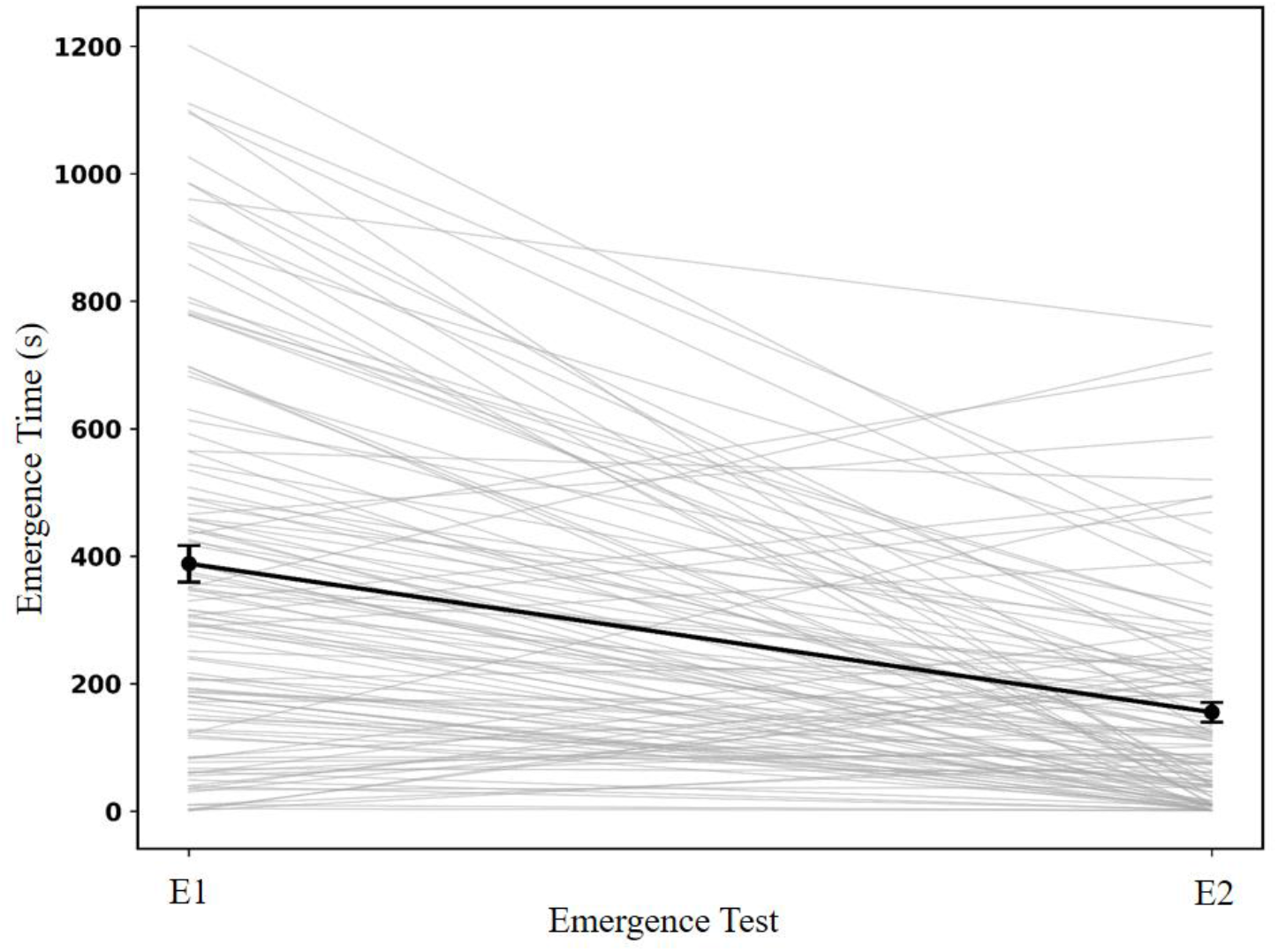
Individual changes in emergence latency between repeated emergence tests. Lines represent individual zebrafish trajectories from the first emergence test (E1) to the second emergence test (E2). Points and error bars represent the mean ± SE emergence latency at each test. The plot illustrates substantial inter-individual variation in emergence latency and the overall change in latency between E1 and E2 (paired t-test: t=8.090, P<0.001).

### 3.3. Association between behavioural phenotype and latency to reach upper half

To further examine whether behavioural phenotype was associated with subsequent locomotor behaviour, latency to reach upper half was compared between bold and shy zebrafish classified according to emergence latency in the first emergence test (E1) (Figure 6). Of the 112 zebrafish, 43 were classified as bold and 28 as shy, whereas Intermediate individuals were excluded from this comparison. One shy individual failed to reach the upper half within the 10 min observation period and was therefore excluded from this analysis. Bold zebrafish showed a lower mean latency to reach upper half (74.00 ± 13.23 s) than shy zebrafish (120.79 ± 24.84 s). However, this difference was not statistically significant (independent t-test, P = 0.104). Considerable inter-individual variation in latency to reach upper half was observed within both behavioural phenotypes.

**Figure 6.**
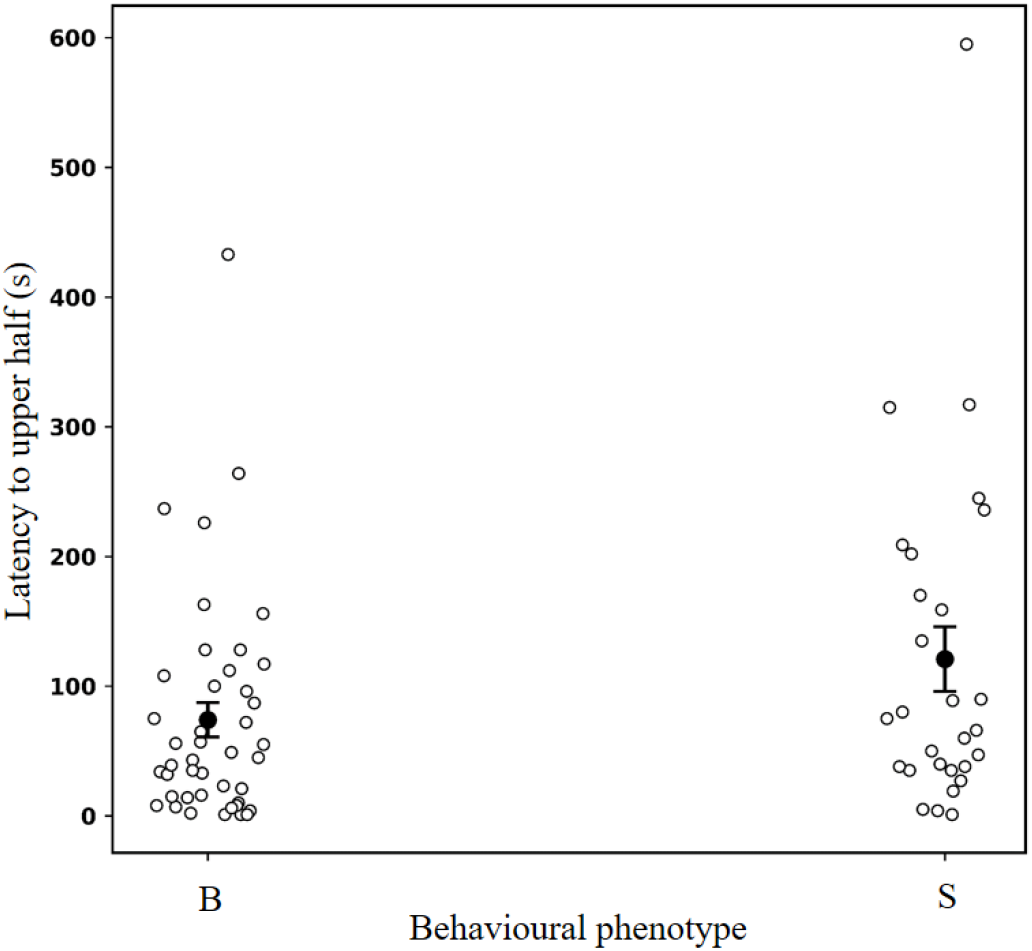
Latency to reach upper half in Bold and Shy zebrafish. Individual latency to reach upper half for zebrafish classified as Bold (B) or Shy (S) according to emergence latency in the first emergence test (E1). Open circles represent individual fish, while filled circles and error bars represent mean ± SE. Intermediate individuals were excluded from the comparison. The difference between behavioural phenotypes was not statistically significant (independent-samples t-test: t_70_ = −1.66, P>0.05).

An exploratory comparison using the E2-based classification also detected no significant difference in upper-half latency between bold and shy fish (independent t-test; t = 0.083, df = 34.12, P = 0.935; Table S1).

### 3.4. Effect of sound exposure on the number of midline crossings

The number of crossings across the horizontal midline of the aquarium (i.e., the boundary separating the upper and lower halves of the water column) was compared between the before (20 s before exposure) and during (20 s exposure) periods for both sound and ambient treatments. In the sound treatment, a significant decrease was observed in the number of midline crossings for the ‘during’ period compared to the ‘before’ period (paired t-test: t_55_=2.835, P<0.01; Figure 7). In contrast, no significant difference was found between the ‘before’ and ‘during’ periods in the ambient treatment (paired t-test: t_55_=-0.645, P>0.05; Figure 7). The change in midline crossings (during − before) differed significantly between treatments, with a greater reduction in the Sound treatment than in the Ambient treatment (mean change ± SE: −1.50 ± 0.53 versus 0.18 ± 0.28 crossings per 20 s; independent-samples Student’s t-test: t_110_ = −2.811, P = 0.0058). Individual crossing counts for ‘before’ and ‘during’ exposure, together with within-fish changes in the sound and ambient treatments, are shown in Figure S1.

**Figure 7.**
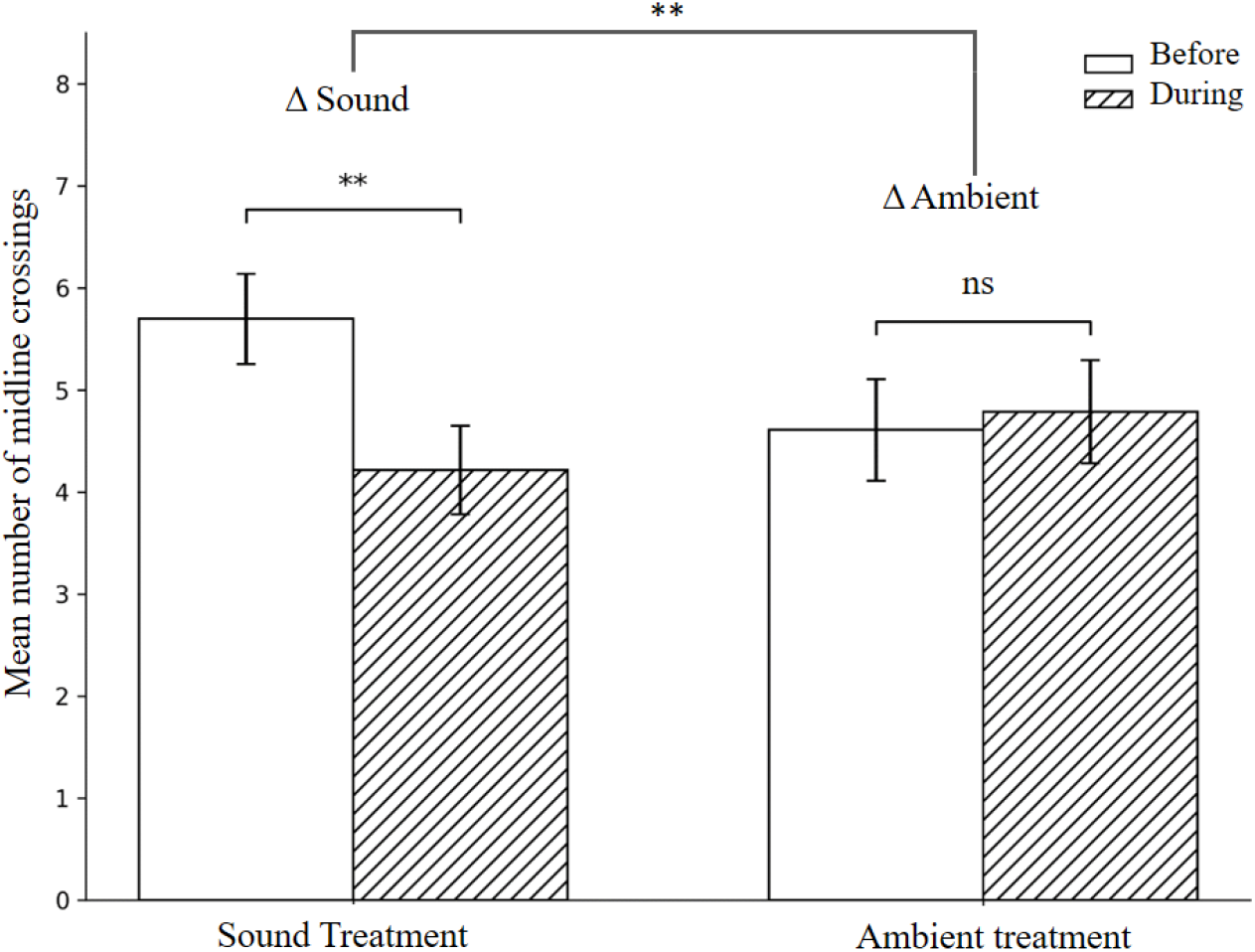
Effect of sound (left panel) and ambient (right panel) treatments on the number of midline crossings per individual (*n* = 56 per treatment). The number of crossings was recorded during the before (20 s before exposure) and during (20 s of exposure) periods. A significant decrease was observed in the Sound treatment (paired t-test: t_55_=2.835, P<0.01), whereas no significant change was detected in the Ambient treatment (paired t-test: t_55_=-0.645, P>0.05). Error bars represent mean ± SEM. ** indicates statistical significance at P < 0.01; ns = not significant (P > 0.05). The between-treatment difference in change scores (during − before) was statistically significant (independent-samples Student’s t-test: t₁ ₁ ₀ = −2.811, P < 0.05).

### 3.5. Correlation between first and second emergence latencies

To assess the relationship between first (ET1) and second (ET2) emergence latencies across repeated trials, we performed a linear regression analysis. The analysis revealed a very weak positive relationship between the two trials (r = 0.263, R² = 0.069, P < 0.01, Figure 8).

**Figure 8.**
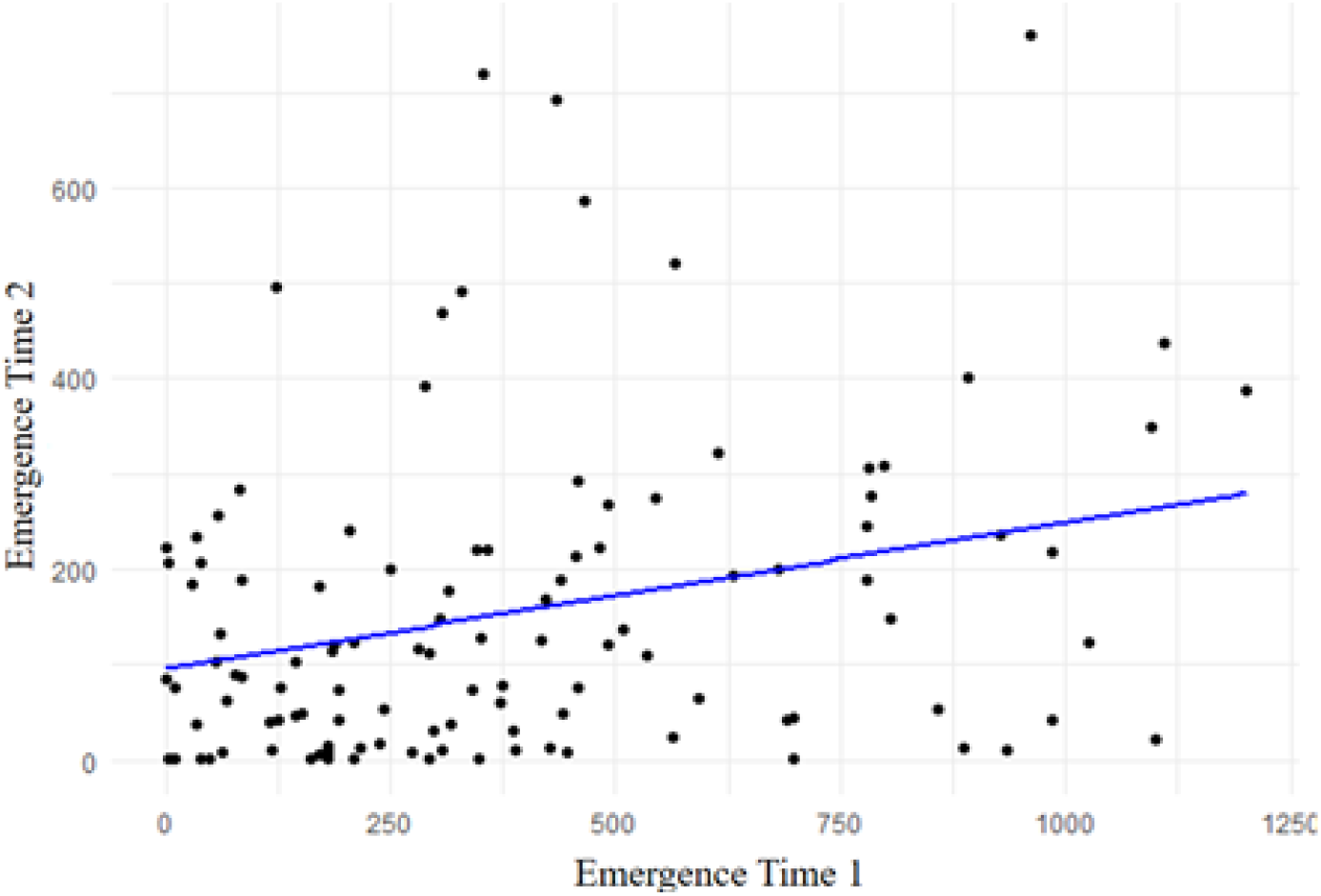
Correlation between first (ET1) and second (ET2) emergence latencies in zebrafish (n=112). The blue line represents the fitted linear regression. Emergence latencies were weakly positively correlated (r = 0.263, R² = 0.069, P < 0.01). This association does not constitute a variance-component estimate of behavioural repeatability.

### 3.6. Association between emergence latency and changes in vertical midline crossings

Exploratory linear models examined whether the association between emergence latency and the change in midline crossings (during − before) differed between the sound and ambient treatments (Figure 9). Using heteroscedasticity-robust HC3 standard errors, the emergence latency × treatment interaction was not statistically significant for either the first emergence test (β = 0.385, 95% CI: −0.043 to 0.813; t_108_ = 1.785, P <0.05) or the second emergence test (β = 0.456, 95% CI: −0.578 to 1.491; t_108_ = 0.874, P < 0.05). Interaction coefficients represent the sound-treatment slope minus the ambient-treatment slope, expressed as the difference in change in crossing counts per 100-s increase in emergence latency. Thus, although the fitted slopes were more positive under sound exposure, there was insufficient evidence that the association between emergence latency and changes in vertical crossings differed between treatments.

**Figure 9.**
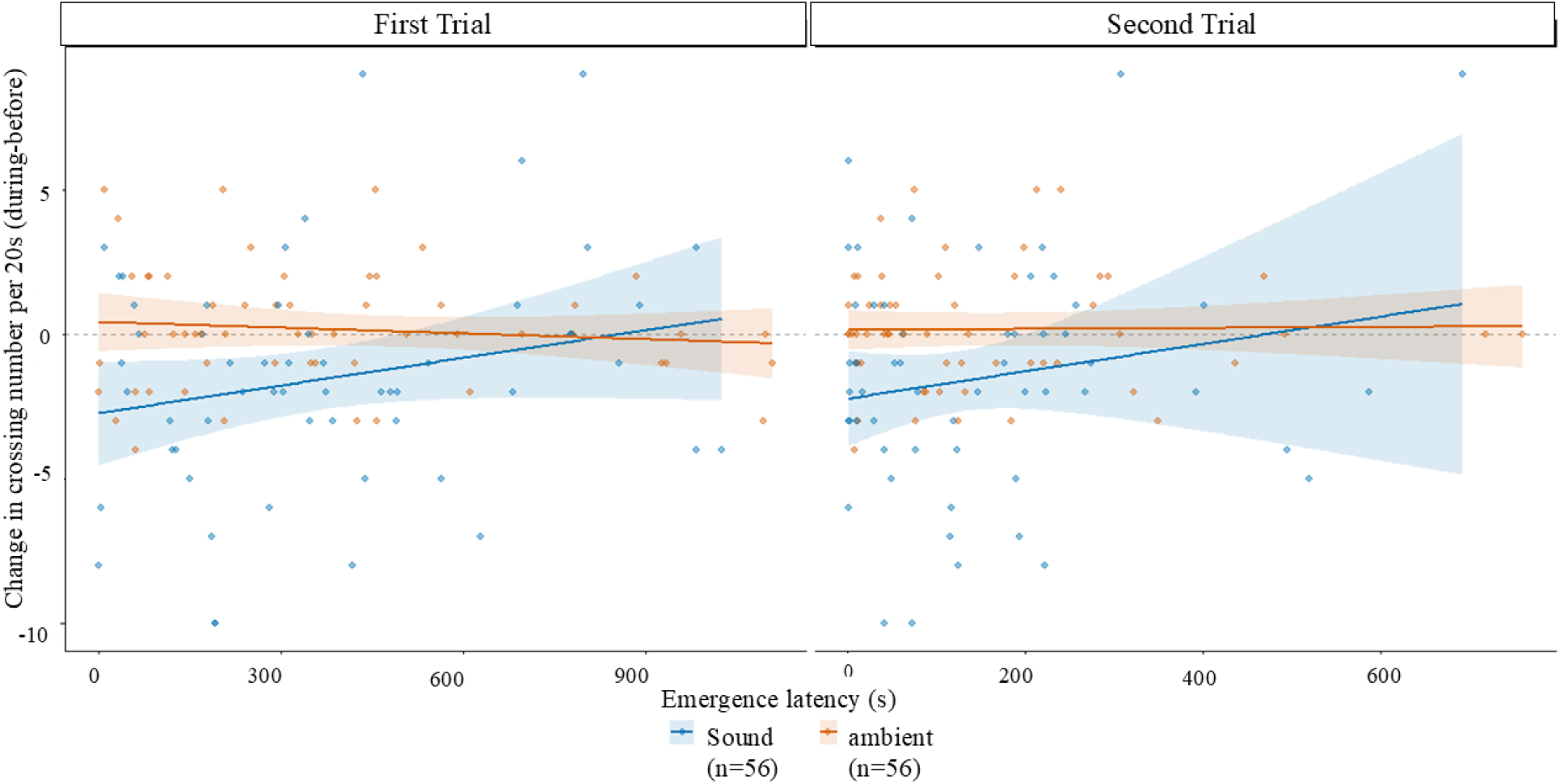
Associations between emergence latency and changes in vertical midline crossings. The left and right panels show the change in crossing counts (during − before) in relation to the first and second emergence latencies, respectively. Both panels use the same fish and the same crossing-change scores; only the emergence-latency predictor differs. Blue and orange points represent the Sound and Ambient treatments, respectively (n = 56 per treatment). Each point represents one fish. Solid lines show treatment-specific linear regression fits, and shaded bands represent 95% confidence intervals for the fitted mean relationships. These bands indicate uncertainty in the estimated relationships, rather than the range of individual behavioural responses.

## 4 Discussion

We investigated whether individual differences in emergence behaviour were associated with changes in vertical movement following sound exposure. Emergence latency decreased between successive tests, while individual latencies were weakly but positively correlated. Upper-half latency did not differ significantly between emergence-based bold and shy categories. Sound exposure caused a higher immediate reduction in vertical midline crossings than ambient conditions. However, the exploratory analyses provided insufficient evidence that the relationship between emergence latency and crossing changes differed between treatments.

### 4.1. Emergence behaviour and individual behavioural variation

Emergence latencies varied among individuals, and fish differed in how much their latencies changed between tests. These observations demonstrate individual variation in emergence behaviour, but do not establish stable personality differences. Because classification thresholds were recalculated for each test, similar proportions of bold, intermediate and shy fish do not reveal whether the same individuals retained their classifications. Assessing consistency requires repeated observations of identified individuals and separation of among-individual differences from within-individual variation (Bell et al., 2009; Dingemanse & Dochtermann., 2013). These categories describe emergence behaviour under the conditions of this experiment and should not be interpreted as evidence of stable personality traits. Using each fish’s emergence latency as a continuous variable allows us to examine whether fish that emerged earlier or later differed in their behavioural responses to sound. Only fish that emerged within the allotted time in both tests were included, so the findings may not generalize to individuals that failed to emerge in either test.

### 4.2. Effects of repeated testing on emergence latency and behavioural consistency

Fish emerged earlier in the second test, while emergence latencies were weakly but significantly positively correlated across tests. Thus, earlier-emerging fish in the first test tended to emerge earlier in the second, despite the overall decrease in latency. This suggests some short-term consistency in relative emergence tendencies, without establishing long-term behavioural stability. The cause of earlier emergence remains unclear because all fish experienced the aquaria in a fixed order, and the second aquarium contained a loudspeaker compartment and mesh barrier. Familiarity, apparatus configuration, and recapture and transfer may all have contributed to the change. Faster emergence alone therefore does not demonstrate learning or habituation; identifying habituation requires distinguishing it from alternative explanations for a declining response (Rankin et al., 2009).

Previous zebrafish research found that activity and exploration remained repeatable over extended periods; although repeatability was lower between the earliest assessments, emphasizing the relevance of prior tank experience (Thomson et al., 2020). Consistent individual differences can therefore coexist with changes in behavioural expression (Dingemanse et al., 2010). Similarly, repeated refuge-emergence tests in wild-type zebrafish showed that behavioural patterns of individual variation responded differently to environmental conditions and prior experience (Singh & Bhat., 2026). Although that study manipulated turbidity rather than sound, it reinforces the distinction between a change in average emergence latency and consistency among individuals across assessments.

### 4.3. Association between emergence-based phenotypes and exploratory behaviour

Emergence latency and upper-half latency describe different stages of behaviour: leaving shelter and subsequently entering the upper water column. Fish classified as bold reached the upper half sooner on average than those classified as shy, but the difference was not statistically significant. This does not establish equivalence between categories (Lakens., 2017), nor does it demonstrate that the two behavioural measures are unrelated. Previous work in zebrafish also indicates that the expression and consistency of emergence-based behavioural profiles depend on the context examined (Mamede et al., 2024).

The comparison included only bold and shy individuals, excluding intermediate responses and reducing the available sample. One shy fish that did not reach the upper half within the observation period was also excluded, so the result applies only to fish that completed this response. Categorisation further removes information about variation within groups (Altman & Royston., 2006). Analysing emergence latency as a continuous predictor would retain this information and allow intermediate individuals to be included. Future analyses should also account explicitly for fish that do not reach the upper half within the observation period.

### 4.4. Behavioural responses during sound and ambient treatments: interpretation and limitations

Vertical midline crossings decreased more in the Sound treatment than in the ambient treatment, supporting an immediate effect of playback on vertical movement. Fewer crossings, however, do not necessarily mean less swimming: fish may have continued moving within one zone. In a multi-laboratory novel tank study, top-entry counts were more strongly associated with distance travelled than with time spent in the upper zone, illustrating the distinction between movement and zone occupancy (Hillman et al., 2025). Previous sound-exposure studies reported changes in zebrafish swimming behaviour and group cohesion without spatial avoidance (Neo et al., 2015a), and a prolonged reduction in swimming speed (Shafiei Sabet et al., 2016). In guppy shoals, noise reduced following events without significantly changing speed or cohesion (Clark et al., 2025). Measuring speed and zone occupancy alongside crossings would help determine whether the present response reflected reduced swimming or a redistribution of movement.

The experimental setting also limits interpretation. Depth and visual conditions influence zebrafish vertical movement behaviour (Blaser & Rosemberg., 2012; Shinkai et al., 2025), so upper-half latency and crossings should be interpreted as movement measures within this shallow arena, rather than validated indicators of anxiety. Equal water depth supports comparisons between treatments, but the findings may not extend to deeper tanks. Particle motion, an important component of sound detection in fishes (Nedelec et al., 2016), was not measured, leaving this aspect of the acoustic exposure uncharacterized. Two-dimensional recordings could not resolve movement towards or away from the camera, limiting conclusions about overall swimming behaviour (Macrì et al., 2017). The short observation windows captured the immediate response to playback onset but did not establish whether it persisted. Crossing counts also provide no direct measure of bottom dwelling, and physiological stress was not assessed. Finally, the genetic background and rearing history of these commercially sourced fish were uncharacterized, so their contributions to individual variation could not be determined.

### 4.5 Emergence latency and individual variation in responses to sound

The exploratory analyses did not provide clear evidence that the relationship between emergence latency and changes in vertical crossings differed between the Sound and Ambient treatments. The present analyses therefore do not establish emergence latency as a predictor of a sound-specific response, although the nonsignificant interactions do not demonstrate equivalent relationships between treatments (Lakens., 2017). In juvenile large yellow croaker, larger changes in behavioural lateralization and performance following boat-noise exposure were reported in fish classified as lateralized before exposure (Xu et al., 2026). Although this finding concerns a different species and trait, it supports further investigation of how initial behavioural characteristics relate to responses to noise. Repeated assessments of emergence and acoustic responses would help separate consistent differences among individuals from variation within individuals (Dingemanse & Dochtermann., 2013). Exposure order also warrants attention: a study involving multiple environmental stressors found that zebrafish responses to noise depended on preceding acoustic conditions (Al Shuraiqi & Barry., 2025).

## 5 Conclusions

Zebrafish emerged sooner in the second test, but the weak positive association between emergence latencies provided only limited evidence of consistency across successive assessments. Sound playback produced a greater immediate reduction in vertical midline crossings than ambient conditions, indicating a change in vertical movement. However, the exploratory analyses did not establish that emergence latency was associated with a sound-specific behavioural response. Repeated assessments of both emergence behaviour and acoustic responses are needed to determine whether individual differences in responsiveness to sound persist over time.

## Acknowledgments

The authors thank the Department of Fisheries, Faculty of Natural Resources, University of Guilan, for providing laboratory facilities and support for this study. We also extend our sincere gratitude to Ms. Zahra Samandar and Ms. Marziyeh Heydari for their invaluable assistance, insightful ideas, and emotional support throughout the course of this research.

## Notes

ChatGPT and DeepL Write were used for language refinement, and ChatGPT also assisted with development of R code for statistical analyses and data visualization. Scite.ai and Google Scholar Labs were used to support literature searches and reference identification. The authors reviewed the text and references and take full responsibility for the content.

## Conflicts of Interest

The authors declare no competing financial interest.

## Notes

### Competing Interest Statement

The authors have declared no competing interest.

